# The effects of human umbilical cord mesenchymal stem cell transplantation on female fertility preservation in mice

**DOI:** 10.1101/2020.03.18.996751

**Authors:** Qiwei Liu, Junhui Zhang, Yong Tang, Yuanyuan Ma, Zhigang Xue, Jinjuan Wang

## Abstract

Female fertility is the capacity to produce oocytes and achieve fertilization and pregnancy, and these outcomes are impaired by age, diseases, environment and social pressure. However, there is no effective therapy that preserves female reproductive ability. Mesenchymal stromal cells (MSCs) can exhibit multidirectional differentiation potential, and they have gained great attention as a tool for preserving female fertility. Therefore, this study uses human umbilical cords-MSCs (Huc-MSCs) to preserve and restore fertility in aging female mice and chemotherapy-damaged mice through the rescue of ovarian function and the reconstruction of the fallopian tubes and uterus. In our study, 2 mouse models were generated: aging mice (37 weeks old) and chemotherapy-damaged mice. Then, we injected Huc-MSCs into mice through the tail vein. After treatment, the effect of MSCs on the ovary, fallopian tubes and uterus was evaluated by analyzing gonadal hormone levels and by performing morphological analysis and statistical analysis. The levels of E2 and FSH exhibited a significant recovery after HUC-MSC transplantation both in aging mice and mice treated with chemotherapy. Huc-MSC treatment also increased the numbers of primordial, developing and preovulatory follicles in the ovaries of mice. Meanwhile, MSCs have been shown to rescue the morphology of the fallopian tubes and uterus through mechanisms such as regenerating the cilia in fallopian tubes and reforming glands and chorionic villi in the uterus. Therefore, it is suggested that Huc-MSCs may represent an effective potential treatment for preserving female fertility through recovery from chemotherapy damage and rescuing female reproductive organs from the effects of aging.

## 1. Introduction

Mesenchymal stromal cells (MSCs) are multipotent adult stem cells that have potent immunomodulatory properties, including the abilities to inhibit the maturation of monocytes, decrease mast cell degranulation, and inhibit B cell proliferation(Shin, et al. 2017). MSCs also can self-renew, have multidirectional differentiation potential, and produce paracrine factors, which have anti-apoptotic, anti-inflammatory and anti-oxidative properties that play a role in restoring injured tissues(Kadam, et al. 2019; Staal, et al. 2011). It has been reported that MSCs release microvesicles containing proteins, mRNA and bioactive lipids, which exert a protective effect on tissues and stimulate tissue repair(Yang, et al. 2019). Because of the above characteristics, MSCs have been successfully used in cell-based therapy for several diseases, such as Asherman’s syndrome, hepatocellular carcinoma, systemic sclerosis and type I diabetes(Liu, et al. 2019; Mohamed, et al. 2019). Recently, MSCs have gained great attention as a tool for preserving female fertility, which could be achieved through recovery of ovarian function and repair of injured uteri(Liu, et al. 2019; Zhang, et al. 2018).

Female fertility is the capacity to produce oocytes to achieve fertilization and pregnancy, and these outcomes are impaired by age, diseases, and environment and social pressure. Age is one of the most important factors that influences female reproductive function(Gumulka and Kapkowska 2005; Silvestris, et al. 2019). Not only does the spontaneous probability of conception start to decrease by 25-30 years of age, but the clinical features of menopausal syndrome also impair the quality of life for women. Menopausal syndrome, which is accompanied by vasomotor phenomena (hot flashes and night sweats) and psychosomatic symptoms (anxiety, feeling of pressure, and depression), is a threat to a women’s physical and mental health. Moreover, the global trend of postponing maternity results in increasing involuntary childlessness mainly because of the age-correlated deterioration of reproductive organs. It has been reported that the number of infertile women with advanced reproductive age (above 40 years old) increased from 9% to 41% from 2003 to 2009 in the USA(Garcia-Velasco, et al. 2013). In China, because of the opening of China’s two-child policy, the population of women becoming pregnant at an advanced reproductive age is increasing. According to a national maternal and child health monitoring report, 17.13% of pregnant women in China were in advanced maternal age in 2017, while this ratio was only 8.4% in 2014. Therefore, maintaining female reproductive ability longer is a current requirement for women.

On the other hand, the incidence of cancer increases dramatically from approximately 1 in 10,000 shortly after birth to approximately 1 in 300 as women reach the end of reproductive life (during their mid-forties)(Bleyer, et al. 2006). Chemotherapy can lead to irreversible female infertility, and nearly 80% of women with cancer face premature ovarian insufficiency (POI) after chemotherapy(Critchley, et al. 2002). POI can cause infertility because of the exhaustion of ovarian follicles. However, there is no effective therapy to preserve female reproductive(Yang, et al. 2019).

In our study, we first aimed to make use of MSCs to repair female reproductive tissue both in aging mice and in chemotherapy-damaged mice. We collected MSCs from human umbilical cords because they are relatively easy to collect, they have few ethical tissue and can be repeatedly obtained. Female reproductive requires the function of the ovary, fallopian tube and uterus, so we have observed the effectiveness of Huc-MSCs in the recovery of those organs to assess, which may become a potential treatment for preserving female productivity.

## 2. Materials and Methods

### 2.1 Isolation and Culture of Huc-MSCs

HUCs were collected from delivering women during uncomplicated caesarean sections after written informed consent was obtained. Participants tested negative for HIV-1, hepatitis B and C, and venereal diseases. HUCs were obtained from Tongji Hospital, Shanghai, China, and the institutional ethics committee approved this project ((TONG) NO.325).

A 10-15 cm piece of HUC was obtained, and the blood was removed. It was washed three times in a penicillin-streptomycin solution (PBS) with penicillin-streptomycin, and then blood vessels were removed on a clean bench. We cut HUCs into approximately 1 mm^3^ pieces. Because MSCs have the ability to migrate from the tissue and adhere to plastic, we spread the HUCs evenly on 100 mm Petri dishes. After 5 min, we added 10 ml of culture medium (89% Dulbecco’s modified Eagle medium: nutrient mixture F-12 (DMEM/F12), 10% fetal bovine serum (FBS) and 1% glutamine (Gln)) to Petri dishes and cultured them directly at 37°C.

After Huc-MSCs were explanted for 7 days, we discarded the culture medium and added 5 ml of new culture medium to resuspend the remaining tissues. Then, the tissue fluid was centrifuged at 1000 rpm for 3 min. Tissues were moved into a new 100 mm Petri dish with 10 ml of culture medium. We obtained MSCs on plastic after culture in a 37°C incubator for 5 days.

### 2.2 Flow Cytometry Analysis

MSCs are defined as having CD44, CD73, CD90, and CD105 upregulated on the cell surface, while MSCs do not express CD45, CD34, CD11b, CD19 and HLA-DR(El Omar, et al. 2014). To assess the immunophenotypes of cultured Huc-MSCs, we performed flow cytometry analysis. Samples were incubated for 30 min in the dark at 4°C after adding relevant antibodies (BD biosciences, Franklin Lakes, NJ). A FACSCalibur system (BD biosciences, Franklin Lakes, NJ) was used to analyze 1×10^4^ cells, and the data were analyzed by FlowJo software.

### 2.3 Growth curve and cycle of cultured MSCs

To test the growth of Huc-MSCs, 5×10^3^ cells/cm^2^ were transferred to 24-well plates on the first day. We counted the number of Huc-MSCs in 3-well plates per day to calculate the average until the 8^th^ day. We used propidium iodide (PI) to stain MSCs; then, they were incubated at 37°C in the dark for 30 min and were finally stored at 4°C in the dark. We performed flow cytometry for red fluorescence at a wavelength of 488 nm, and we analyzed the cell scattering.

### 2.4 Biological activity of Huc-MSCs

To assess the biological activity of Huc-MSCs, we induced cell differentiation into chondrocytes, adipocytes and osteocytes separately. Huc-MSCs (5×10^3^ cells/cm^2^) were explanted onto cell slides, and then the culture media removed and differentiation culture medium was added when cell growth reached 60%. Cells were incubated at 37°C. All differentiation kits were purchased from Life Technologies. After inducing differentiation for 25 days, we stained cells with alcian blue stain, green neutral lipid stain and alizarin red s stain and observed cells using a microscope.

### 2.5 Animals and Huc-MSC transplantation

Female C57 mice aged 6-7 weeks were fed under standard laboratory conditions (14 h of light, 10 h of dark; 25°C). Sixty female mice were divided into two groups: 20 mice were in the control group, and 40 mice were in the experimental group. Mice from the experimental group (CTx-mice) were administered busulfan (20 mg/kg, #2635, Fluka) and cyclophosphamide (200 mg/kg, #C0768, Sigma) by intraperitoneal injection; each drug was dissolved in dimethyl sulfoxide (DMSO) (#D2650, Sigma). Mice from the control group (non-CTx-mice) were injected with a volume of DMSO that matched the volume injected into the experimental group. One week after chemotherapy, 2×10^6^ Huc-MSCs in 50 μl of PBS at the 7th passage were transplanted into 20 CTx-mice through the tail vein. Twenty CTx-mice and 20 non-CTx-mice were injected with only PBS.

Alternatively, there were 20 female mice, aged 38 weeks, that were divided into two groups: an experimental group was transplanted with 2×106 Huc-MSCs and a control group was transplanted only with PBS.

### 2.6 Enzyme-linked immunosorbent (ELISA) assay

Mouse blood was extracted from the caudal vein of mice and placed at room temperature for 2 h. Then, mouse blood was centrifuged at 2000 g for 20 min, and the supernatant was collected. The E2 and FSH levels were measured using an ELISA kit (Cloud-Clone Corp, USA) according to the manufacturer’s protocol, and the results were analyzed using a Synergy H4 Hybrid Reader.

### 2.7 Morphological Analysis and Ovarian Follicle Counting

Ovaries, fallopian tubes and uteri were dissected from mice 2, 4 and 6 weeks after the Huc-MSC injection. Briefly, the tissues were fixed with 10% formaldehyde solution for at least 24 h. Tissues were dehydrated with alcohol, and then the whole tissues were placed in paraffin and stored in the incubator. Then paraffin-embedded tissues were sectioned, the slide were dried at 45°C and hematoxylin (H) and eosin (E) were used to stain the slides. Last, the slides were dehydrated and fixed. Tissue histological examination was performed under light microscopy (Nikon ECLIPSE 80i, Japan). The follicles were classified as primordial, primary, secondary and early antral follicles and were counted.

### 2.8 Immunofluorescence staining

To detect Huc-MSCs in the ovary, ovarian sections were immunostained with a CD74 antibody (dilution 1:100) and were incubated overnight at 4°C. Then, the sections were washed 3 times and were incubated with goat anti-rabbit IgG secondary antibody for 60 min at 37°C. Then, the nuclei were stained with DAPI and washed 4 times with PBS. Last, slides were incubated with an anti-fluorescence quencher.

### 2.9 Statistical analyses

All continuous data were performed by using SPSS 22.0 (IBM SPSS Statistics for Macintosh, Version 22.0 Armonk, NY: IBM Corp.). Continuous variables were described as the mean±SD, and t-tests were used for continuous variables. A two-sided P-value <0.05 was considered significant.

## 3. Results

### 3.1 Characterization of Huc-MSCs and location in the ovary

Flow cytometric analysis showed that cells strongly expressed CD44, CD73, CD90, and CD105 but not CD45, CD34, CD11b, CD19 or HLA-DR (Figure 1A). We also tested cell growth at different passages (Figure 1C). It was shown that cell growth was relatively stable at different passages. Most cells stayed at G0 and G1 stage during the proliferation process (Figure 2A). Huc-MSCs were induced to form osteocytes after culturing in osteogenic medium (Figure 2B). After culturing in chondrogenic medium, Huc-MSCs were induced to form chondrocytes (Figure 2C). Culturing in adipogenic medium induced Huc-MSCs to form adipocytes (Figure 2D). Huc-MSCs were observed in the ovarian tissue by labeling with a CD 74 antibody after they were injected into the mouse tail vein (Figure 3). Above all, those outcomes revealed that these cells were MSCs and had the characteristics of stem cells.

**Figure 1.**
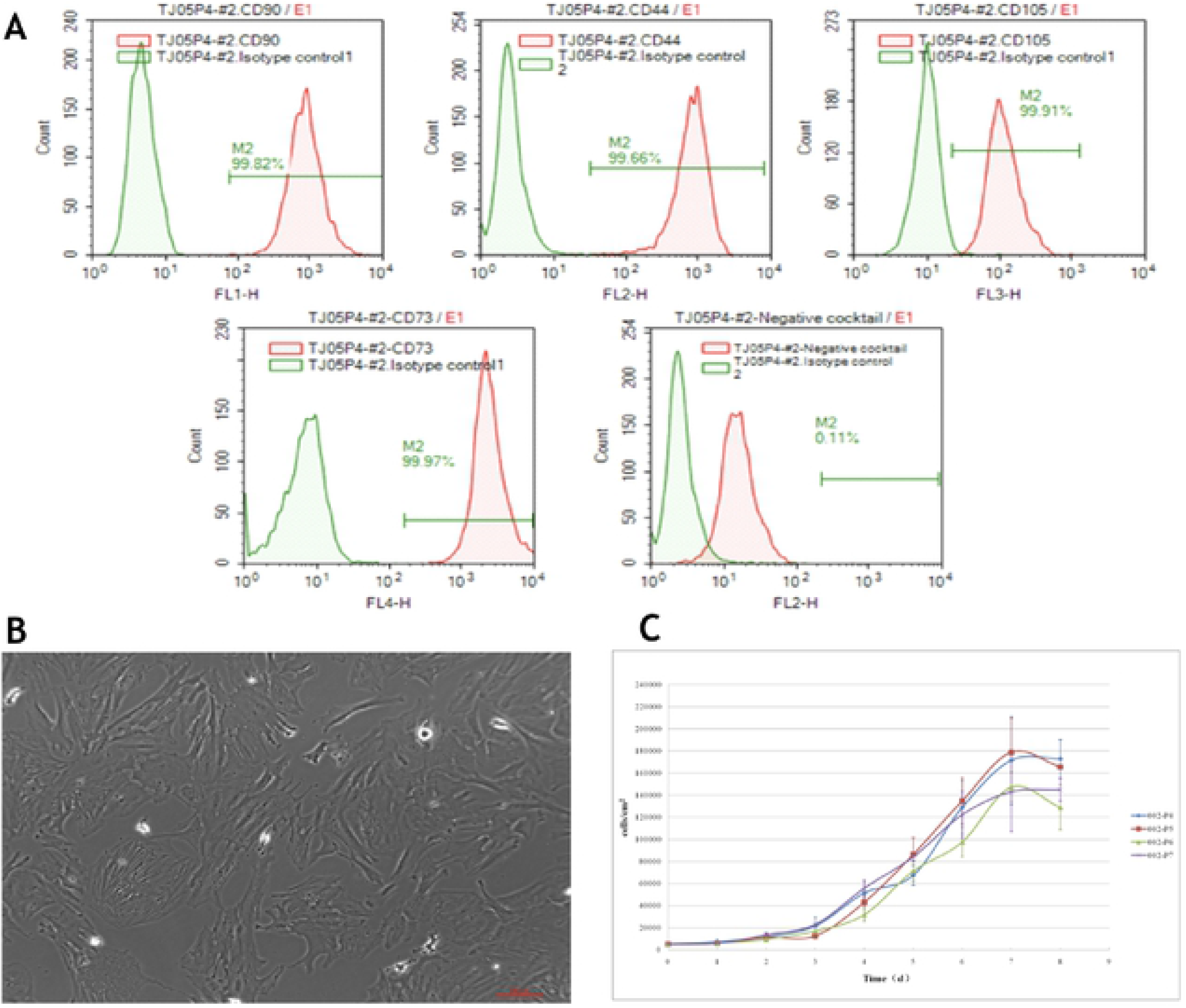
Characterizations of Huc-MSCs in culture. Immunophenotypes of human umbilical cord stem cells expressed markers of MSCs (A). Huc-MSCs exhibit fibroblast-like morphology and adhere to the surface of plastic cell culture dishes (B). Growth curve of Huc-MSCs at fifth generation (C).

**Figure 2.**
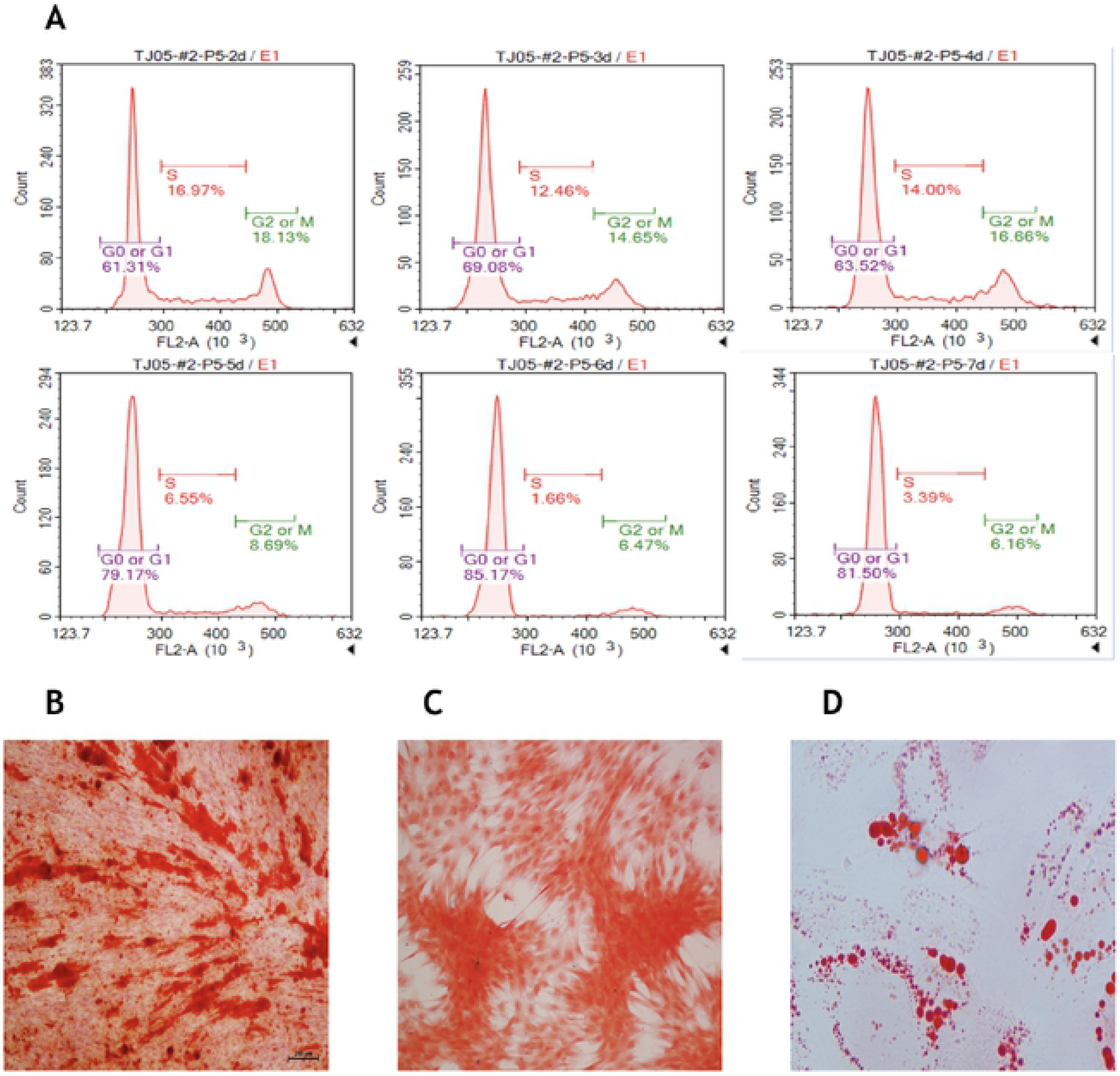
Detecting the cell cycle and induction of Huc-MSCs. Detecting the cell cycles of MSCs culturing from day 3 to day 7 (**A**). Cell viability is best during day 3 and day 5. Huc-MSCs can be induced into chondroblast (**B**), osteoblasts (**C**) and adipocyte (**D**).

**Figure 3.**
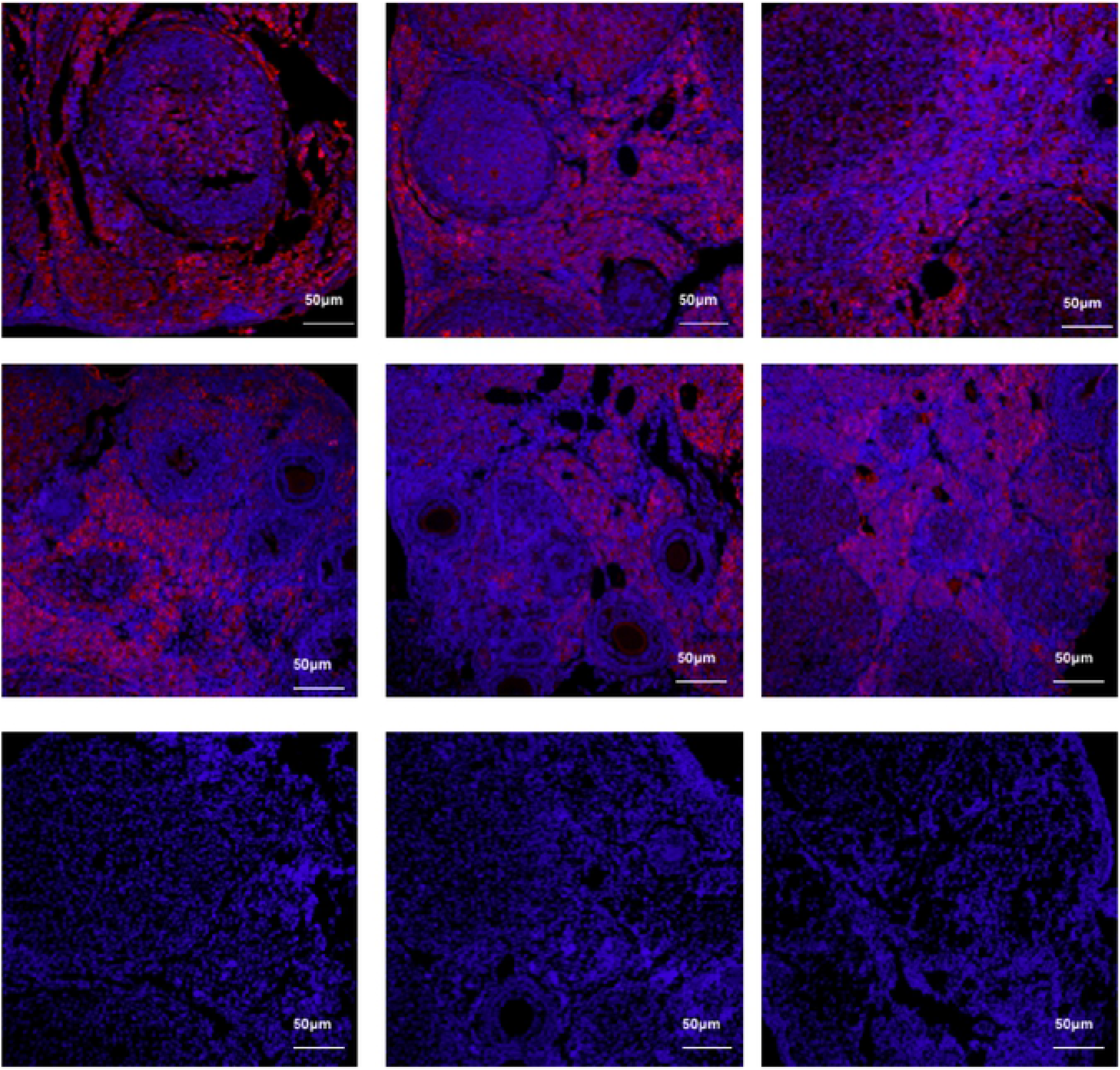
Localization of Huc-MSCs in mouse ovaries. The nuclei were stained with DAPI (blue), and Huc-MSCs were stained with red. After Huc-MSCs transplantation, Huc-MSCs can be traced in mice ovaries.

### 3.2 Huc-MSCs improve sex hormone secretion in chemotherapy-damaged and aging ovaries

Two, four and six weeks after Huc-MSC transplantation, levels of sex hormones, including E2 and FSH, were detected. The serum levels of E2 were significantly higher, and the levels of FSH were significantly lower in POI mice with Huc-MSCs injection when compared with the levels in POI mice without MSC treatment (P < 0.05) (Figure 4-1, 4-2). The same results also showed that elderly mice with Huc-MSCs expressed higher E2 and lower FSH when compared to the levels in aging mice without Huc-MSCs (P < 0.05) (Figure 4-3, 4-4).

**Figure 4.**
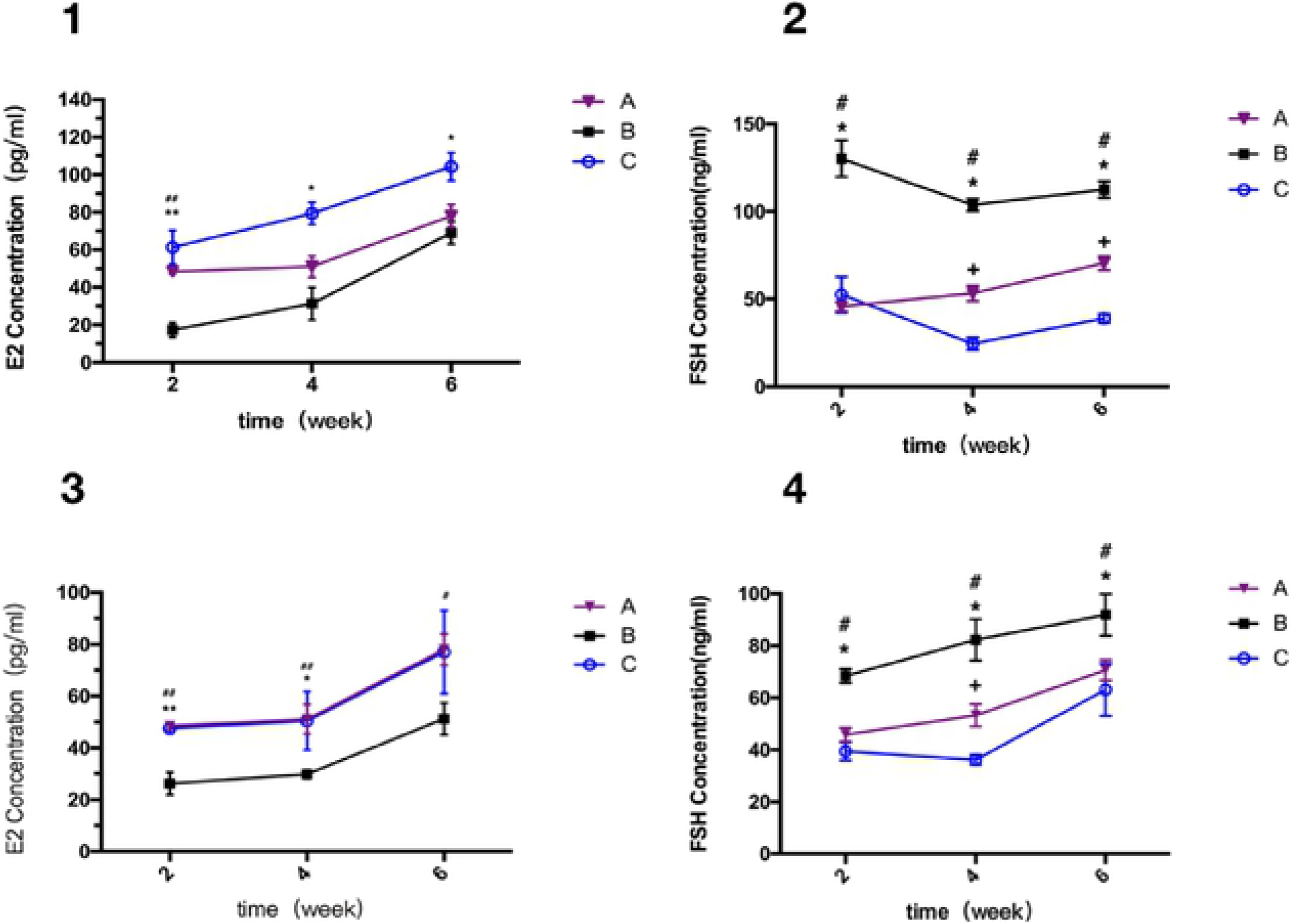
Effects of Huc-MSC transplantation on serum hormone release in mice. 1, 2: A: control mice; B: chemotherapy-damaged (CTx) mice without MSCs; C: CTx mice with MSCs. 3,4: A: control mice: B: aging mice without MSCs; C: aging mice with MSCs. *: B versus C has statistical significance (*P*<0.05); #: A versus B has statistical significance (*P*<0.05); +: A versus C has statistical significance (*P*<0.05).

### 3.3 Huc-MSCs stimulate oocyte production in aging mice

In our study, we injected Huc-MSCs into the mouse tail vein, and their ovaries were collected and observed at 2 weeks, 4 weeks and 6 weeks after injection. For the aged mice experiment, ovaries from 37-, 39- and 41-week-old mice had the appearance of an atrophic cortex with loss of follicles (Figure 5A, B, C). The cortex of the ovary in aging mice was filled with degenerated follicles that were either cystic or contained only several granular cells. However, with Huc-MSC treatment, the ovaries of aging mice were improved (Figure 5D, E, F). At 2 and 4 weeks of treatment, ovarian folliculogenesis was restored. The number of total follicles decreased in aging mice over time. However, after Huc-MSC treatment, the level of follicles reached the level of the control mice at 7 weeks of age (Figure 5G-K). It can be seen that there were obviously increasing numbers of secondary and antral follicles.

**Figure 5.**
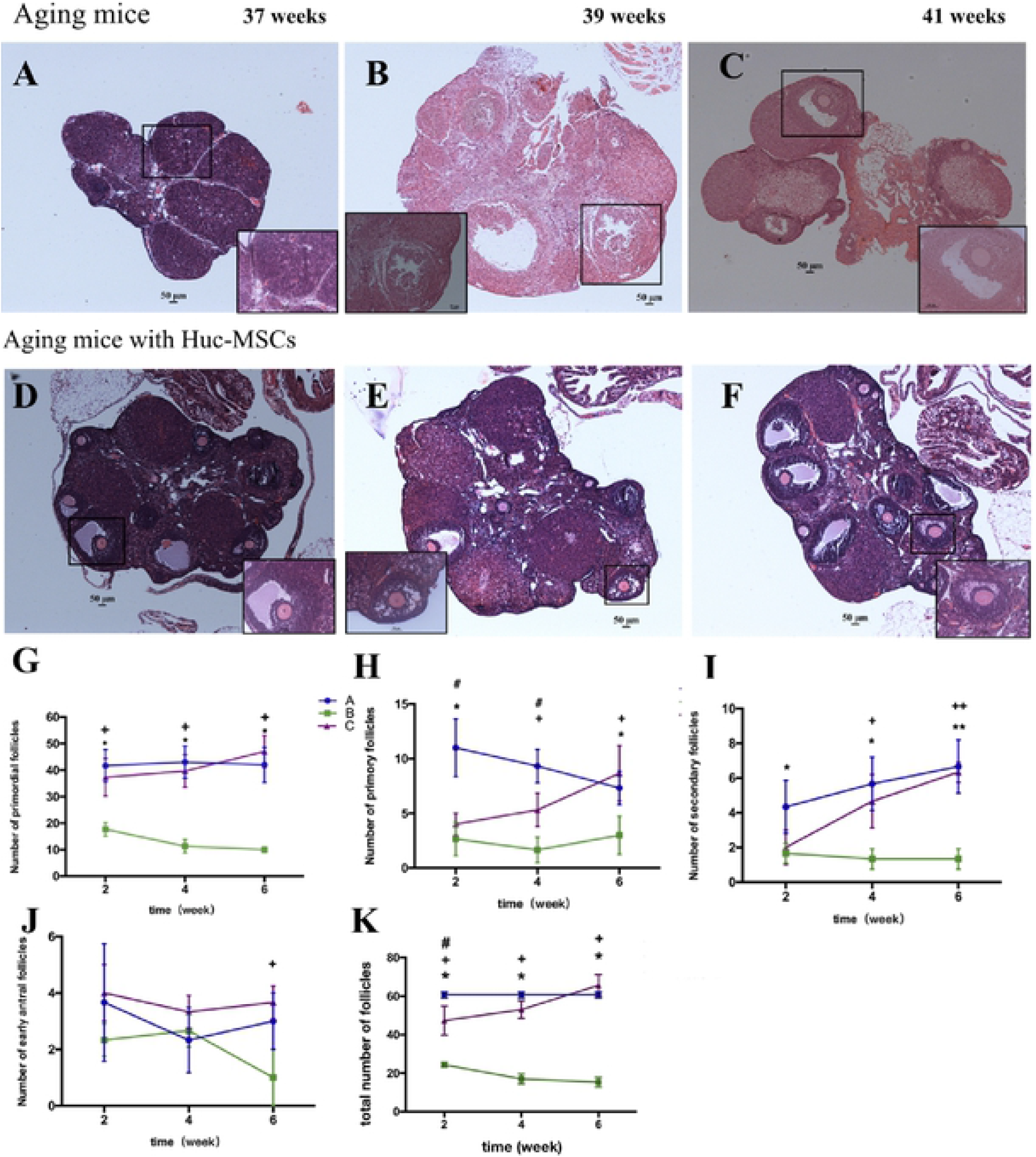
Effects of Huc-MSC transplantation on ovarian morphology and summary of ovarian follicle numbers in each stage of aging mice. A-C: Aging mice without Huc-MSC treatment. A: 37-week-old mice without Huc-MSC treatment. B: 39-week-old mice without Huc-MSC treatment. C: 41-week-old mice without Huc-MSC treatment. D-F: Aging mice treated with Huc-MSCs. D: 37-week-old mice treated with Huc-MSCs. F: 39-week-old mice treated with Huc-MSCs. F:41-week-old mice treated with Huc-MSCs. G-J: Summary of ovarian follicle numbers at each stage, including primordial, primary, secondary and early antral follicles. In figures G-K, A is the control mouse group, B is the aging moue group and C is the aging mouse group with Huc-MSC injections. *: A versus B has statistical significance (*P*<0.05); +: B versus C has statistical significance (*P*<0.05); #: A versus C has statistical significance (*P*<0.05).

### 3.4 Huc-MSC repair tubes and uterus in aging mice

The epithelium of the fallopian tube is ciliated. Destroying cilia or ciliary motion has been proven to cause infertility(Briceag, et al. 2015). The normal fallopian tube had a thin muscular layer and spindly and regular cilia (Figure 6A). Figure 6B shows that the inner surface of the tubes lined by ciliated cells was damaged, mucosal folds disappeared, and the lumen of the uterine tube was also narrowed, which may have caused a loss of ciliary activity. However, after MSC treatment, cilia were restored and appeared to be spindly and located regularly around the epithelium of the fallopian tube (Figure 6C). For the 38-week uterus, endometrial glands were rarely observed, and chorionic villi had disappeared (Figure 6E). After injection of Huc-MSCs, the construction of the endometrium was obviously improved (Figure 6F).

**Figure 6.**
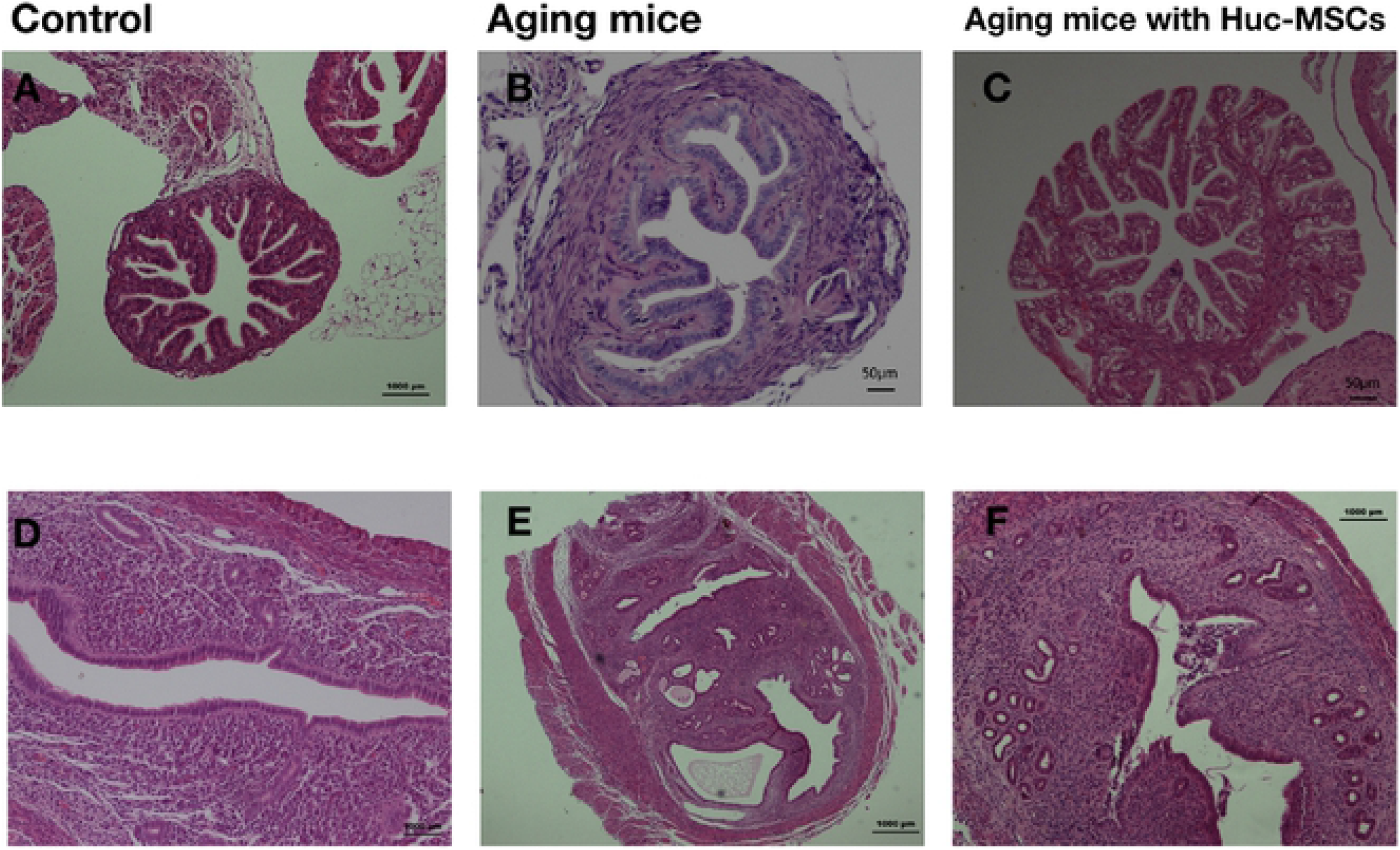
Effects of Huc-MSC transplantation on fallopian tubes and uterus morphology in aging mice. A-C is fallopian tubes. A: wild mice at 7 weeks. B: 39-week-old mice. C: 39-week-old mice with Huc-MSCs. D-F is uterus. D: wild mice at 7 weeks. E: 39-week-old mice. F: 39-week-old mice with Huc-MSCs.

### 3.5 Huc-MSCs stimulate oocyte production in chemotherapy-damaged mice

Figure 7A, D and G show ovaries of a normal size from control mice at 9, 11 and 13 weeks of age, respectively; these ovaries were composed of variant stage follicles, including primordial, primary, secondary and antral follicles. In Figures 5B, E and H, the number of follicles decreased steeply in chemotherapy-damaged mice that did not receive an Huc-MSC injection at ages of 9, 11 and 13 weeks (Figure 7B, E, H). Follicles were deteriorated, most of which even exhibited a loss of oocytes in the intracavity. These figures also showed multiple stages of follicles after chemotherapy was stopped. However, compared with POI mice without treatment, the numbers of follicles were greater in the mice that received the Huc-MSC injection (Figures 7C, F and I). Furthermore, follicles in ovaries with Huc-MSC treatment were structurally sound and were present in different stages. From Figure 7J-N, it can be clearly seen that with the treatment of Huc-MSCs, the numbers of follicles in the treatment group at various stages were all greater than that of the chemotherapy-damaged mice without treatment (P<0.05). However, the number of follicles in the experimental group did not reach the level of the normal group.

**Figure 7.**
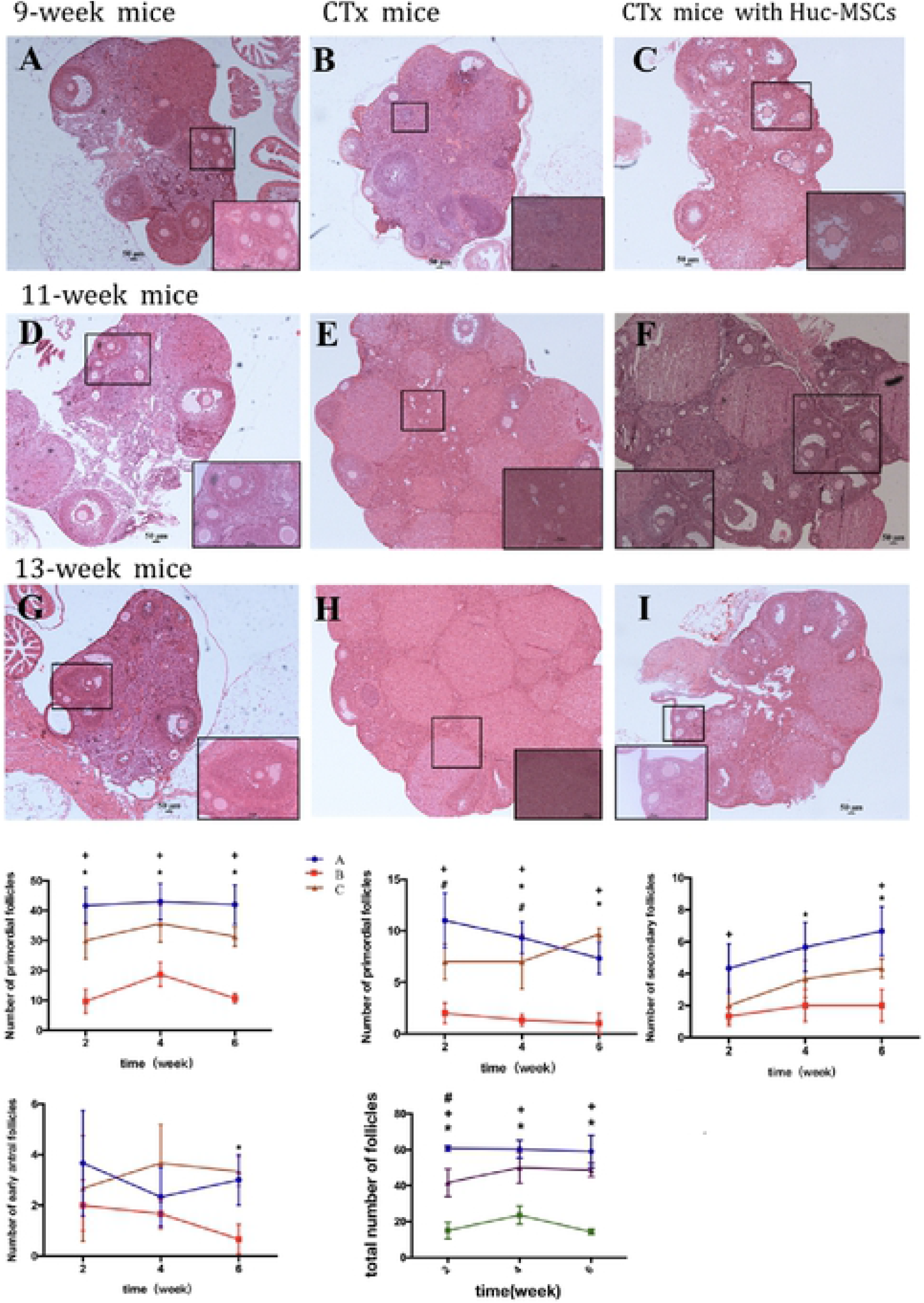
Effects of Huc-MSC transplantation on ovarian morphology and summary of ovarian follicle numbers in each stage. A, D, G: wild mice at 9 weeks, 11 weeks and 13 weeks respectively. B, E, F: chemotherapy-damaged (CT) mice without Huc-MSCs treatment at 9 weeks, 11 weeks and 13 weeks respectively. C, F I: CT mice with Huc-MSCs treatment at 9 weeks, 11 weeks and 13 weeks respectively. J-N: summary of ovarian follicles at each stage, including primordial, primary, secondary and early antral follicles. In J-N figures, A is wild mice group. B is CT mice group without Huc-MSCs injections. C is CT mice group with Huc-MSCs injections. *: A versus B has statistical significance (*P*<0.05); +: B versus C has statistical significance (*P*<0.05); #: A versus C has statistical significance (*P*<0.05).

### 3.6 Huc-MSCs repair the construction of fallopian tubes and uterus in chemotherapy-damaged mice

Chemotherapy damaged the fallopian tube structure, as shown in Figure 8B; the cavity of the tube was shrunken, and the cilia disappeared. However, 6 weeks after treatment, the cilia of the fallopian tube had formed, the muscular layer became thinner and the cavity expanded (Figure 8C).

**Figure 8.**
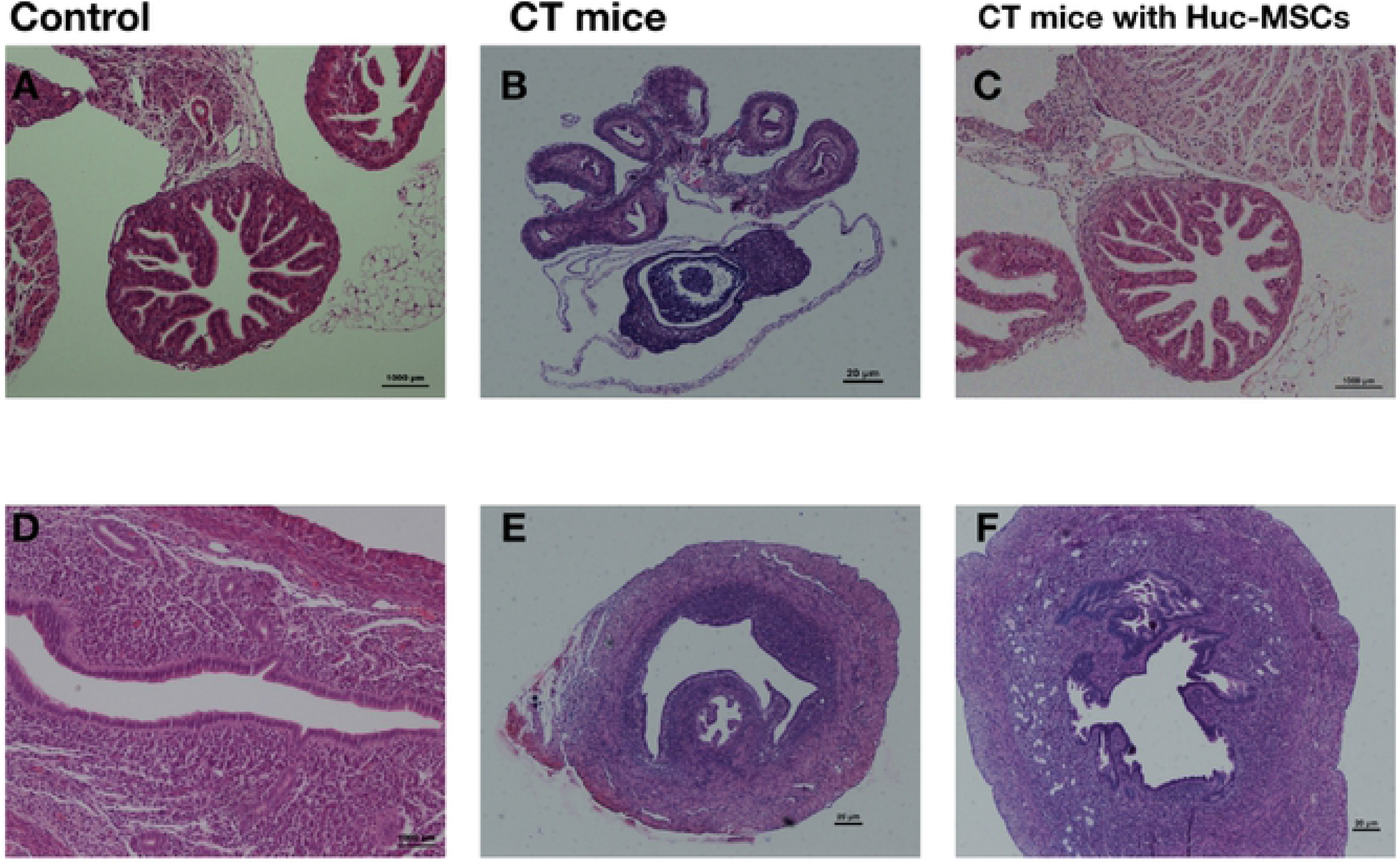
Effects of Huc-MSCs transplantation on fallopian tubes and uterus morphology in mice. A-C is fallopian tubes. A: wild mice at 7 weeks. B: 9-week-old mice without Huc-MSCs treatment. C: 9-week-old mice with Huc-MSCs treatment. D-F is uterus. D: wild mice at 7 weeks. E: 9-week-old mice without Huc-MSCs treatment. F: 9-week-old mice with Huc-MSCs treatment.

The healthy uterine endometrium was composed of glands and chorionic villi. After chemotherapy, endometrial tissue with rare glands became thinner, and chorionic villi disappeared (Figure 8E). The mesometrium was loose, and cells composing the serosal layer were irregular. The uterus was improved after Huc-MSC injection. Chorionic villi were formed, and endometrial villi with many glands were restored and proliferated (Figure 8F). It was also clearly shown that the mesometrium and serosal layer were dense and well distributed.

## 4. Discussion

In our study, we focused on two main factors that decrease female fertility, chemotherapy and natural aging. Our study is the first to make full use of Huc-MSCs in the preservation and recovery of fertility in female aging mice and chemotherapy-damaged mice; the recovery included rescuing ovarian function and reconstruction of the fallopian tubes and uterus. The positive effect of Huc-MSCs in infertility treatment was confirmed in our study. We observed that the number of follicles in chemotherapy-damaged mice improved dramatically after Huc-MSC treatment. Second, the numbers of primordial, developing and preovulatory follicles in the ovaries of both POI and aged mice were significantly higher than that of the control group. Meanwhile, our results also show the disturbed estrous cycle of POI and aging mice. The levels of E2 and FSH exhibited a significant recovery after HUC-MSC transplantation, which was a response to the recovery of ovarian function. Moreover, we have also found that Huc-MSCs could effectively improve the morphology of the fallopian tubes and uterus, by performing act such as regenerating the cilia in fallopian tubes and reforming the glands and chorionic villi in the uterus. Therefore, it is suggested that Huc-MSCs can not only improve ovarian function but also induce reconstruction of the fallopian tubes and uterus in mice.

Age is the single most important factor in determining fertility potential in females, and it is supposed to be an irreversible process. In recent years, childbearing has been increasingly delayed, which leads to an increased demand for preserving fertility as long as possible. Aging women more easily develop complete ovarian failure because of their low primordial follicle pool(Brice, et al. 2002; Rivkees and Crawford 1988). Meanwhile, as increasing nuclear genome abnormalities occur with age, the quality of oocytes appears to decline(May-Panloup, et al. 2016). It is suggested that in women older than 38 years, the luteinizing granulosa cells are less numerous, there are lower concentrations of sex hormones and there is increased mitochondrial damage(Vollenhoven and Hunt 2018). The impact of maternal age on the uterus also decreases the success of pregnancy(Woods, et al. 2017). This may be because the aging uterus cannot provide a normal placental and uterine environment for embryonic development. A previous study showed that in older female mice, the number of live offspring declined significantly even when the number of early implantation sites was the same as that of young mice(Woods, et al. 2017). Age is also an important factor causing tubal factor infertility^(Maheshwari, et al. 2008)^. This experiment reported that the ratio of tubal factor infertility in women over 35 years old is significantly higher than it is in women under 30 years old. Second, chemotherapy damages the function of the ovary mainly because it can induce the apoptosis of granulosa cells, it can damage the ovarian microenvironment, and it can injure growing follicles(Roness, et al. 2014; Song, et al. 2016). Chemotherapy also damaged the morphology of fallopian tubes, as shown in our study.

Based on the capacity of multilineage development, relatively easy differentiation, immunomodulation, and long-term ex vivo culture, MSCs are becoming attractive cells in clinical treatment of neuronal impairment, functional recovery of the brain and treatment myocardial injury(Gutierrez-Fernandez, et al. 2013; Rangappa, et al. 2003; Yang, et al. 2015). In 2005, the Johnson group found that bone marrow transplanted into the ovary can restore oocytes in mice following chemotherapy treatment(Johnson, et al. 2005). Since then, to treat female infertility, researchers have used MSCs from various tissues, such as fat tissue, endometrium, amniotic fluid stem cells, placenta and umbilical cord blood(Bianco, et al. 2008; Jin, et al. 2016; Lai, et al. 2015; Xiao, et al. 2014), which provide new methods for preserving female fertility. Several studies have reported that MSCs can be differentiated into granulosa cells and reduce apoptosis of granulosa cells so that MSCs can repair ovarian function for women after chemotherapy. MSCs can also provide a variety of cytokines and chemokines, including epidermal growth factor (EGF), insulin-like growth factor binding protein (IGFBP), and insulin-like growth factor-1 (IGF), for target cells through paracrine processes(Islam, et al. 2016; Zhang, et al. 2016). Several studies have suggested that MSCs stimulate the expression of anti-inflammatory cytokines, such as IL10 and tumor necrosis factor (TNF)-α, which improve the function of damaged fallopian tubes by decreasing the level of inflammatory factors(Li, et al. 2017; Liao, et al. 2019). These secretomes have shown immense potential for the treatment of damaged endometrium by promoting endometrial proliferation and angiogenesis.

## 5. Conclusion

In our study, the ovarian function was recovered in chemotherapy-damaged mice and aging mice by Huc-MSC treatment, and the morphology of the endometrium and fallopian tubes were also obviously improved. Above all, Huc-MSCs may be an effective potential treatment for preserving female reproductive function, by recovering after chemotherapy damage and rescuing female productive organs from the effects of aging.

## 6. Acknowledgment

This research was supported by grant of Beijing Obstetrics and Gynecology Hospital, Capital Medical University (FCYY201813).

## 7. Author Contribution

XZG and JJW designed the study. LQW, ZJH and TY built mice models, cultured Huc-MSCs, and did ELISA assay.MYY finished morphological analysis and statistical analyses. LQW and ZJH made charts. All authors read and approved the final manuscript.

